# Causal Evidence for Lateral Prefrontal Cortex Dynamics Supporting Cognitive Control

**DOI:** 10.1101/130096

**Authors:** Derek Evan Nee, Mark D’Esposito

## Abstract

The lateral prefrontal cortex (LPFC) is essential for higher-level cognition, but how interactions among LPFC areas support cognitive control has remained elusive. In previous work, dynamic causal modeling (DCM) of fMRI data revealed that demands on cognitive control elicited a convergence of influences towards mid LPFC. We proposed that these findings reflect the integration of abstract, rostral and concrete, caudal influences to inform context-appropriate action. Here, we provide a causal test of this model using continuous theta-burst transcranial magnetic stimulation (cTBS). cTBS was applied to caudal, mid, or rostral LPFC, as well as a control site in counterbalanced sessions. In most cases, behavioral modulations resulting from cTBS could be predicted based upon the direction of influences within the previously estimated DCM. However, inconsistent with our DCM, we found that cTBS to caudal LPFC impaired cognitive control processes presumed to involve rostral LPFC. Revising the original DCM with a pathway from caudal LPFC to rostral LPFC significantly improved the fitted DCM and accounted for the observed behavioral findings. These data provide causal evidence for LPFC dynamics supporting cognitive control and demonstrate the utility of combining DCM with causal manipulations to create, test, and refine models of cognition.

## Introduction

Context-appropriate behavior requires assessing both present circumstances and future plans to determine the best course of action in the moment. Guiding behavior in accordance with internal representations rather than habitual stimulus-response tendencies requires cognitive control which depends on the lateral prefrontal cortex (LPFC; Miller and Cohen, 2001). Yet, how the functional properties and interactions among areas of the LPFC support cognitive control remains poorly understood.

Previously, we collected fMRI data on human participants to investigate how the LPFC supports cognitive control by examining intercommunication among LPFC areas across a variety of cognitive control demands (Figure 1A,B; Nee and D’Esposito, 2016). First, we used univariate analysis to determine the functional responses of different LPFC areas. This analysis revealed that caudal LPFC was responsive to attention to stimulus features (*Feature Control*), mid LPFC was responsive to contextual rules to be applied to the attended stimulus feature (*Contextual Control*), and rostral LPFC was responsive to retaining a past stimulus feature for a future rule (*Temporal Control*). Collectively, these results supported the hypothesis that progressively rostral LPFC areas perform progressively abstract processes (Fuster, 2001; Koechlin et al., 2003; Badre, 2008; Badre and D’Esposito, 2009). Second, we used dynamic causal modeling (DCM) to examine how intercommunication among LPFC areas supports cognitive control (Friston et al., 2003). The estimated model (Figure 1D) demonstrated three principle properties: 1) caudal LPFC provides feature-specific inputs to the rest of the LPFC; 2) rostral and caudal influences converge in mid LPFC during *Contextual Control*; and 3) rostral LPFC remains largely segregated from the rest of the LPFC during *Temporal Control* (i.e. functional modulations do not extend to the rest of LPFC; Figure 1D, red arrows). Collectively, these results indicated that the mid LPFC is a nexus where multiple influences converge to guide context-appropriate action, providing a new framework upon which to understand how the functional interactions within the LPFC support cognitive control.

**Figure 1:**
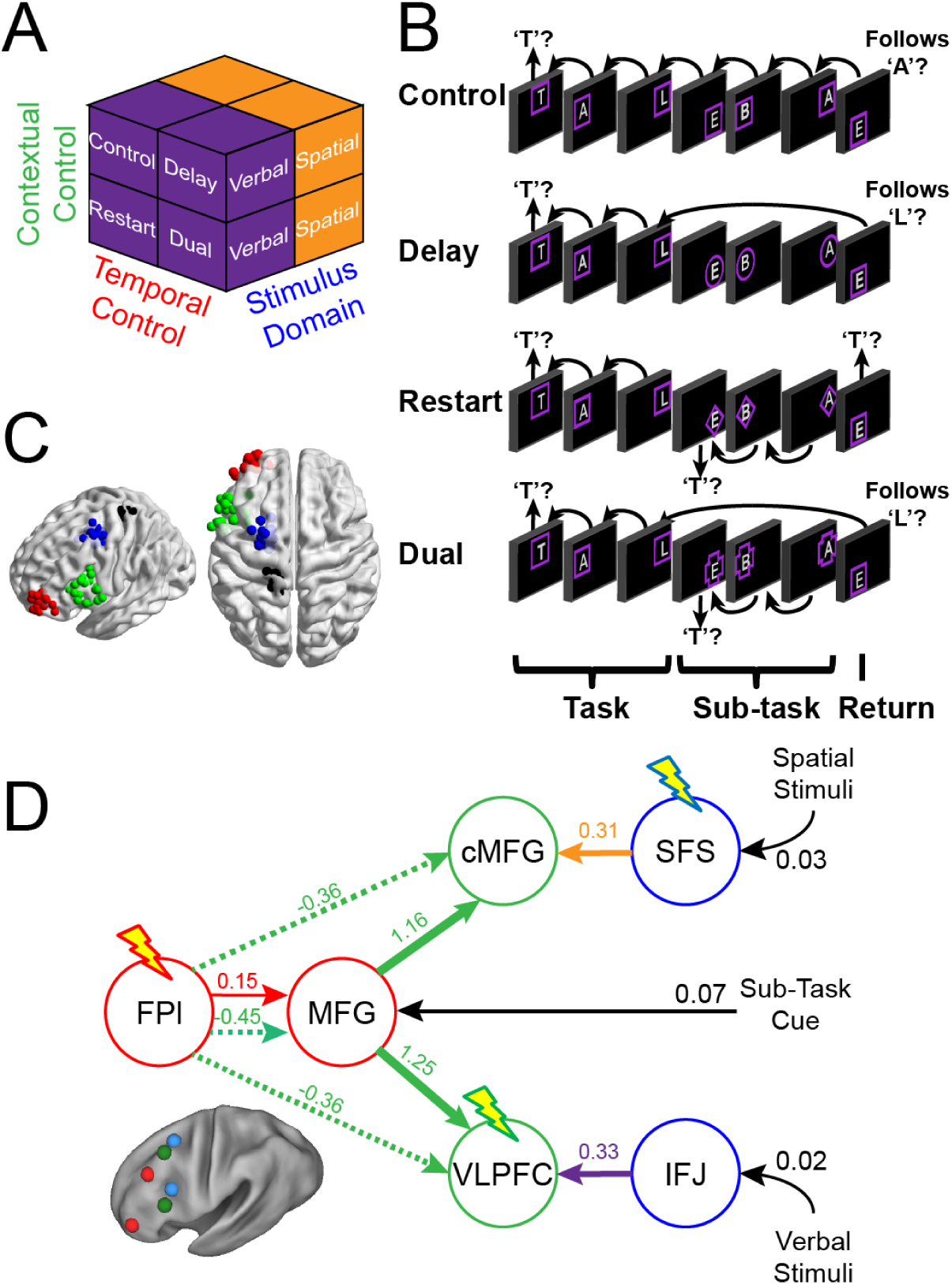
Task, cTBS targets, and dynamic causal model. A) The design orthogonally manipulated factors of *Stimulus Domain* (verbal, spatial) and two forms of cognitive control: *Contextual Control* (low – *Control*, *Delay*; high – *Restart*, *Dual*) and *Temporal Control* (low – *Control*, *Restart*; high – *Delay*, *Dual*). These factors were fully crossed in a 2 × 2 × 2 design. B) The basic task required participants to judge whether a stimulus followed the previous stimulus in a sequence. The sequence in the verbal task was the order of the letters in the word “tablet.” The sequence in the spatial task was a trace of the points of a star. The start of each sequence began with a decision regarding whether the currently viewed stimulus is the start of the sequence (e.g. ‘t’ in the verbal task). Factors were blocked with each block containing a basic task phase, a sub-task phase, a return trial, and a second basic task phase (not depicted), for all but the *Control* blocks. *Control* blocks consisted only of the basic task phase extended to match the other conditions in duration. Colored frames indicated whether letters or locations were relevant for the block (verbal – purple; spatial – orange in this example; verbal condition depicted). The basic task was cued by square frames. Other frames cued the different sub-task conditions. In the *Delay* condition (circle frames), participants held in mind the place in the sequence across a distractor-filled delay. In the *Restart* condition (diamond frames), participants started a new sequence. In the *Dual* condition (cross frames), participants simultaneously started a new sequence, and maintained the place in the previous sequence. C) Each sphere represents a stimulus target for an individual. Red – rostral LPFC, green – mid LPFC, blue – caudal LPFC, black – S1 (control site). D) The dynamic causal model estimated previously (Nee and D’Esposito, 2016). Colored arrows denote modulations of effective connectivity during different cognitive control demands (orange – attention to spatial features; purple – attention to verbal features; green – *Contextual Control*; red – *Temporal Control*). Colors of the circles denote univariate sensitivities to *Feature Control* (blue), *Contextual Control* (green), and *Temporal Control* (red). Lightning bolts indicate targets for continuous theta-burst transcranial magnetic stimulation (cTBS). cTBS was predicted to affect behavior for which a given region was responsive, and also behaviors supported by downstream regions that require processing in upstream targets. Abbreviations: SFS – superior frontal sulcus; IFJ – inferior frontal junction; cMFG – caudal middle frontal gyrus; VLPFC – ventrolateral prefrontal cortex; MFG – middle frontal gyrus; FPl – lateral frontal pole.

A central goal of neuroscience is to determine causal associations among stimuli, neural systems, and behavior as a gateway to specifying predictive models of behavior. While DCM has been demonstrated to accurately detail functional neural circuitry (Lee et al., 2006; David et al., 2008; Bernal-Casas et al., 2017) modeling causal links among neural regions, it does not specify the causal relationship between neural interactions and behavior. However, given the complexities of neural interactions, a model of directed influences is a critical intermediary to determine the link between brain and behavior (Jazayeri and Afraz, 2017). That is, a model of directed influences affords predictions regarding how perturbations of specific brain regions affects other regions and ultimately behavior. Within this framework, one can identify parent and children nodes wherein perturbations of parents affects children, but not vice versa. This sort of framework is critical to establish neural hierarchies (Badre et al., 2009; Azuar et al., 2014), and navigate the complex pathways by which neural activity leads to behavior.

Here, we apply this logic by using continuous theta-burst transcranial magnetic stimulation (cTBS) to reversibly reduce cortical excitability (Huang et al., 2005). We apply cTBS to either caudal (superior frontal sulcus; SFS), mid (ventrolateral prefrontal cortex; VLPFC), or rostral LPFC (lateral frontal pole; FPl), as well as a control site (primary somatosensory cortex; S1) in a within-subjects counterbalanced design (Figure 1C). Following the previously estimated DCM, we predicted that 1) cTBS to caudal LPFC would result in a *feature-specific* impairment acting generally across cognitive control demands. This follows from the model prediction that caudal LPFC provides feature inputs to the rest of the LPFC system. 2) cTBS to mid LPFC would result in a *feature-specific* impairment during *Contextual Control*. This follows from the model prediction that mid LPFC integrates feature information from caudal LPFC and task information from mid-rostral LPFC to perform *Contextual Control*. 3) cTBS to rostral LPFC would result in a *feature-general* impairment during *Temporal Control*. This follows from the model prediction that the rostral LPFC is functionally segregated from other LPFC areas during *Temporal Control* and that this region is insensitive to feature information. These patterns of results would provide causal evidence linking the estimated neural dynamics to their presumed behavioral correlates.

## Results

### Replication of Previous fMRI Results

Prior to receiving cTBS, all participants underwent an fMRI session using the same task as previously described (Figure 1A,B; Nee and D’Esposito, 2016) in order to localize individual targets for cTBS (Figure 1C). The new sample offered an opportunity to replicate the previous findings. Each individual performed a single fMRI session in the present study compared to two sessions in the previous report, so the effects reported here are expected to have reduced power relative to the original study. Nevertheless, as depicted in Figure 2 and its supplements, virtually all of the previously reported effects were replicated (replication statistics reported in Figure captions). As before, a progression of activation from caudal to rostral LPFC was observed for *Feature Control* to *Contextual Control* to *Temporal Control* (Figure 2). Sensitivity to stimulus features was present in caudal and mid, but not rostral LPFC consistent with more abstract processing in rostral LPFC (Figure 2 – figure supplement 1). Caudal LPFC was related to present, but not future behavior, rostral LPFC showed the inverse pattern, and mid LPFC showed sensitivity to both present and future behavior, collectively forming a temporal abstraction gradient (Figure 2 – figure supplement 2). Finally, the effective connectivity parameters estimated by DCM were similar to the previous report. The major difference from the previously reported DCM results was that the pathways linking caudal to mid LPFC were not modulated by attention to stimulus features (i.e. *Stimulus Domain*). Instead, the stimulus inputs to caudal LPFC were increased several fold (Figure 2 – figure supplement 3). Intuitively, these different findings reflect whether an attention-related gain is realized prior to (input parameter) or after (modulation of caudal to mid LPFC effective connectivity) feature information arriving at LPFC. Effectively, this leads to the same result – feature information is propagated through the LPFC via input nodes in feature-specific caudal LPFC during attention to a given feature. Statistically, these different findings reflect an attentional gain that is sustained (modulation) versus transient (input). These differences do not affect the predictions of the model with regard to cTBS since either way, disruption of caudal LPFC is predicted to disrupt feature-specific inputs to the rest of the LPFC.

**Figure 2:**
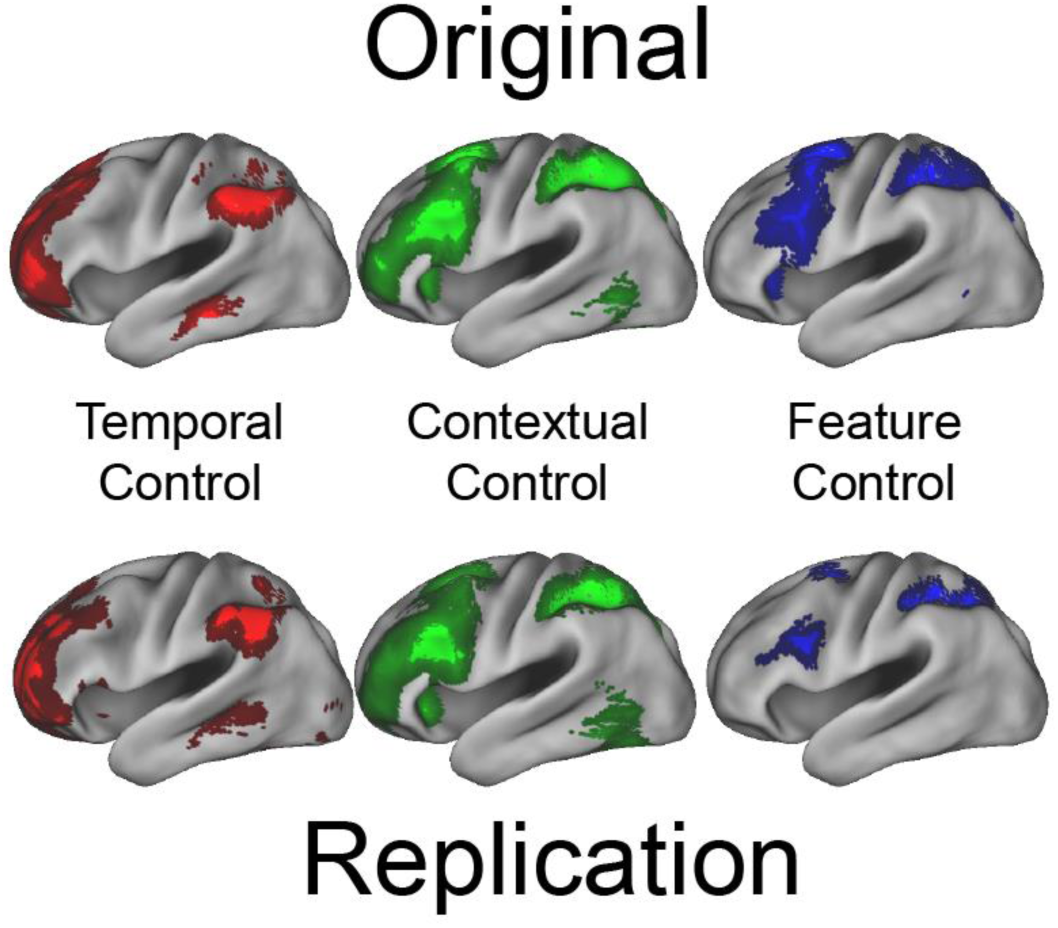
Univariate replication. Top: previously reported univariate effects. Bottom: univariate effects in the present study. In both cases, *Temporal Control* elicited activation of rostral LPFC, *Contextual Control* of mid LPFC, and *Feature Control* of caudal LPFC indicating a rostral-caudal gradient of cognitive control. All results thresholded at p < 0.001 at the voxel level with 166 cluster extent providing family-wise error correction at p < 0.05.

As before, additional analyses on the effective connectivity parameters estimated by DCM were performed to examine hierarchical control and its relationship to higher-level cognitive ability as indexed by a combination of short-term/working memory capacity and fluid intelligence (see Nee and D’Esposito, 2016 for complete details). Replicating the previous results, hierarchical strength, as measured by greater efferent relative to afferent fixed connectivity, peaked in mid-rostral LPFC and was low in FPl, the rostral-most area, suggestive that the rostral-most LPFC is not the apex of the hierarchy (Figure 2 – figure supplement 4). Once again, individuals varied in the strength of bottom-up versus top-down modulations as a function of cognitive control with a tradeoff between the two reflected in a negative correlation between these parameters (Figure 2 – figure supplement 5A). However, whereas we had previously reported that individual differences in top-down modulations as a function of cognitive control (Figure 2 – figure supplement 5B), and hierarchical strength in fixed connectivity (Figure 2 – figure supplement 5D) were both positively related to individual differences in trait-measured higher-level cognitive ability, this relationship was not evident in the present sample. While a null effect might be expected given the reduced power in the present sample relative to the previous one, the trends that were evident were opposite in sign to those observed previously. As reported in more detail below, the positive relationship between DCM parameters and trait-measured higher-level cognitive ability were also not replicated in the revised DCM. Given that nearly all of the other effects replicated, this particular lack of replication may indicate that our previous result was a false positive.

### Behavioral Results and Effects of cTBS

To examine replication of the previously reported behavioral effects of cognitive control demands, separate 2 × 2 × 2 ANOVAs on error-rate (ER), and reaction times (RT) on correct trials were performed for data collected during the fMRI session during the sub-task phase (Figure 3). As before, these analyses revealed significant effects of cognitive control demands with main effects of *Temporal Control* (ER: F(1,23) = 6.04, p < 0.05; RT: F(1,23) = 11.34, p < 0.005), *Contextual Control* (ER: F(1,23) = 60.30, p < 10^-7^; RT: F(1,23) = 149.48, p < 10^-10^), and their interaction (ER: F(1,23) = 6.92, p < 0.05; RT: F(1,23) = 105.30, p < 10^-9^). While no main effect of *Stimulus Domain* was observed previously, in the present sample there was a main effect of *Stimulus Domain* in RT (F(1,23) = 4.28, p < 0.05) but not ER (F(1,23) = 2.33, p > 0.1). Participants performed better on the spatial task relative to the verbal task, an effect which was larger with increased practice as is evident by an increase in the effect size during the cTBS sessions as reported below. Once again, there were no interactions between *Stimulus Domain* and cognitive control demands (ER and RT all p > 0.25), although this also changed with practice as detailed below.

**Figure 3:**
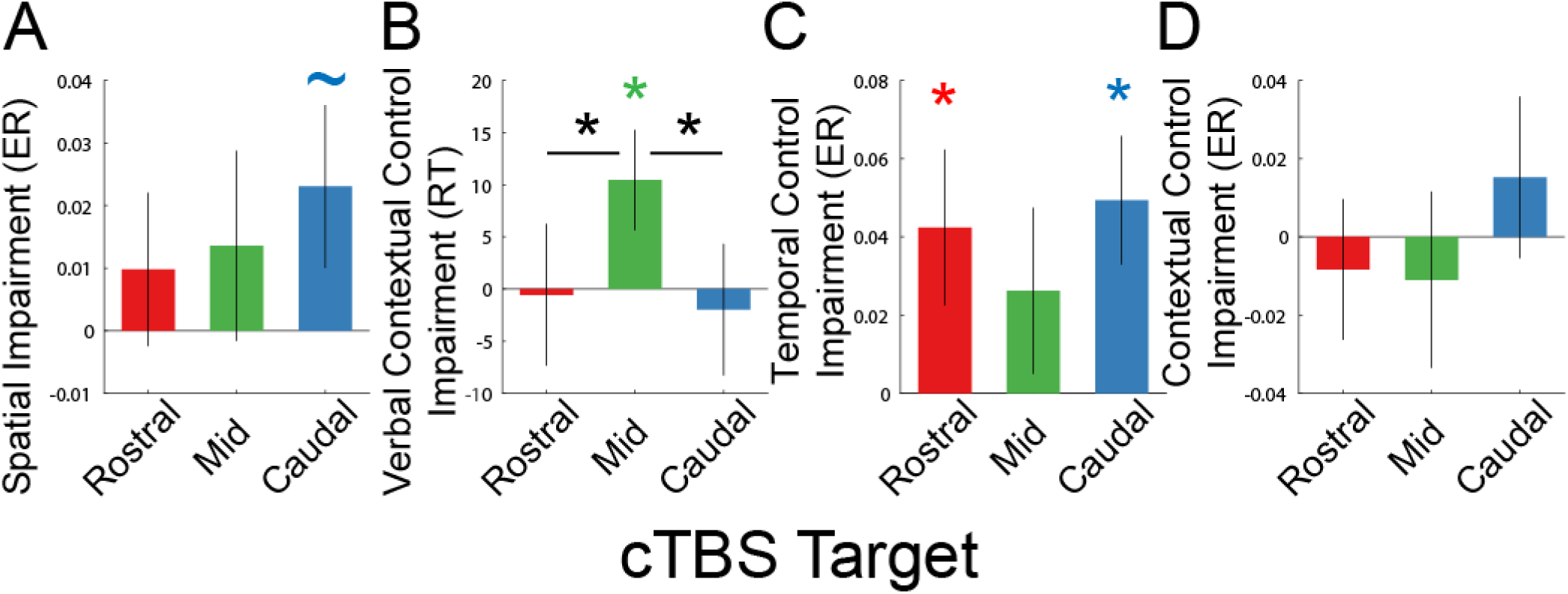
Behavioral effects following cTBS. In all bars, performance following cTBS to the control target (S1) has been subtracted out such that the y-axis indicates a behavioral impairment relative to control stimulation. The x-axis indicates the stimulation target (rostral – rostral LPFC/FPl; mid – mid LPFC/VLPFC; caudal – caudal LPFC/SFS). A) Impairments in *Spatial Stimulus Domain* relative to *Verbal Stimulus Domain* in error-rate (ER). B) The interaction between *Stimulus Domain* × *Contextual Control* in reaction time (RT) contrasted such that relative verbal impairments are positive (i.e. greater verbal *Contextual Control* cost relative to spatial *Contextual Control* cost). C) Impairments in the main effect of *Temporal Control* in ER. D) Impairments in the main effect of *Contextual Control* in ER.

Next, we examined effects of cTBS on behavior. Given that the hypotheses are borne out of interactions that can be difficult to visualize, we depict contrasts that address model predictions in Figure 3. The full uncontrasted data are depicted in Figure 3 – figure supplement 1.

First, the model predicts that cTBS to caudal LPFC would result in a feature-specific impairment given that caudal LPFC provides feature inputs to the rest of the LPFC. In this case, we targeted the caudal superior frontal sulcus (SFS), roughly corresponding to the frontal eye-fields, that was modeled as the source of spatial feature inputs. In this case, we expected diminished spatial relative to verbal task performance to result from cTBS to SFS. To test this prediction, we examined how the effect of *Stimulus Domain* was modulated by cTBS *Target*. A 2 × 4 ANOVA with factors of *Stimulus Domain* (verbal, spatial) and *Target* (FPl, VLPFC, SFS, S1) revealed a significant main effect of *Stimulus Domain* in ER and RT (ER: F(1,22) = 13.75, p < 0.005; RT: F(1,22) = 24.43, p < 0.005), but no main effects of *Target* nor interactions between *Target* and *Stimulus Domain* (ER and RT all p > 0.3). However, a planned contrast comparing cTBS to SFS and S1 revealed a significant *Stimulus Domain* × *Target* interaction in ER (F(1,22) = 3.13, p < 0.05, one-tailed; Figure 3A), but not RT (F(1,22) < 1). While participants generally performed better on the spatial relative to verbal task after cTBS (FPl: t(22) = 3.67, p < 0.005; VLPFC: t(22) = 2.43, p < 0.05; S1: t(22) = 3.04, p < 0.01), this was not the case following cTBS to SFS (t(22) = 1.41, p > 0.15). Hence, cTBS to SFS resulted in a specific impairment in spatial task performance in line with model predictions.

Second, the model predicts that cTBS to mid LPFC would result in a feature-specific impairment in *Contextual Control* given that *Contextual Control* requires integration of bottom-up feature information (from caudal LPFC) and top-down task information (from mid-rostral LPFC). In this case, we targeted the ventrolateral prefrontal cortex (VLPFC), which was sensitive to *Contextual Control* for verbal information. Correspondingly, we expected diminished verbal relative to spatial *Contextual Control* performance to result from cTBS to VLPFC. To test this prediction, we examined whether the *Stimulus Domain* × *Contextual Control* interaction was modulated by cTBS *Target*. A 2 × 2 × 4 ANOVA with factors of *Stimulus Domain*, *Contextual Control*, and *Target* revealed main effects of *Stimulus Domain* (ER: F(1,22) = 6.83, p < 0.05; RT: F(1,22) = 23.14, p < 0.0005) and *Contextual Control* (ER: F(1,22) = 135.22, p < 10^-9^; RT: F(1,22) = 213.97, p < 10^-11^), as well as a *Stimulus Domain* × *Contextual Control* interaction in RT (F(1,22) = 17.30, p < 0.0005). The interaction was driven by reduced costs associated with *Contextual Control* for the spatial relative to verbal task. No main effect of *Target* nor interactions with *Target* were observed (ER and RT all p > 0.1). However, a planned contrast comparing cTBS to VLPFC and S1 revealed a significant *Stimulus Domain* × *Contextual Control* × *Target* interaction in RT (F(1,21) = 4.72, p < 0.05; Figure 3B), but not ER (F(1,21) = 2.51, p > 0.1). This interaction was driven by increased *Contextual Control* costs for the verbal relative to spatial task following cTBS to VLPFC. Furthermore, this same interaction was observed when comparing cTBS to VLPFC with the other LPFC sites (VLPFC vs. FPl: F(1,22) = 4.51, p < 0.05; VLPFC vs. SFS: F(1,22) = 5.87, p < 0.05). Thus, cTBS to VLPFC resulted in a feature-specific, control-specific impairment consistent with predictions of the model.

Finally, we examined the impact of cTBS on the rostral LPFC by targeting FPl. According to the model, effective connectivity from the FPl to mid LPFC is reduced during *Contextual Control*. In the previous report, we suggested that this negative modulation effectively nullifies positive associations between the FPl and mid LPFC in fixed connectivity, thereby serving to segregate the FPl from the rest of the LPFC. If so, cTBS to FPl would not be expected to have an effect on *Contextual Control* since cTBS is expected to perform a similar segregating function by way of reducing FPI cortical excitability. Another possibility is that the negative modulation reflects top-down inhibition from FPl to mid LPFC which may serve to prioritize which contextual information is appropriate at which time. In this case, cTBS to FPl would be expected to affect *Contextual Control* since reducing FPI cortical excitability would, in turn, reduce the top-down bias it transmits. Either way, an effect on *Temporal Control* was anticipated given the region’s sensitivity to *Temporal Control*. To test these predictions, we performed a 2 × 2 × 4 ANOVA with factors of *Contextual Control*, *Temporal Control*, and *Target*. This analysis revealed a main effect of *Contextual Control* (reported above), but no main effects of *Temporal Control* or *Target* (ER and RT, all p > 0.05). *Temporal Control* and *Contextual Control* interacted (ER: F(1,21) = 31.26, p < 0.0001; RT: F(1,21) = 133.69, p < 10^-9^). While there were no significant 2- or 3-way interactions with *Target*, the *Temporal Control* × *Target* interaction approached significance in ER (F(1,21) = 2.47, p = 0.07). Planned contrasts comparing cTBS to FPl and S1 revealed a significant *Temporal Control* × *Target* interaction in ER (F(1,21) = 4.55, p < 0.05; Figure 3C), but not RT (F(1,21) = 1.13, p > 0.3), and no interactions between *Contextual Control* × *Target* either alone (Figure 3D) or with *Temporal Control* (ER and RT, all p > 0.1). Hence, these data indicate that the FPl is causally related to *Temporal Control*, but do not provide evidence that cTBS to FPl affects *Contextual Control*.

Collectively, the analyses above suggest that caudal, mid, and rostral LPFC support separable cognitive control processes, the nature of which is predicted by a model of LPFC dynamics. To further corroborate these associations, we performed a 4 × 3 ANOVA with factors of *Target* and *Contrast* using the three contrasts reported above that were predicted to demonstrate dissociative effects (*Stimulus Domain*, *Stimulus Domain × Contextual Control*, *Temporal Control*). This analysis confirmed a significant interaction (F(6,126) = 2.93, p < 0.05). A similar analysis extended to include the *Contextual Control* contrast was also significant (F(9,189) = 2.13, p < 0.05). Although we cannot conclude a triple dissociation since not all pairwise differences were significant for each measure, this result adds credence to associations of each region to its corresponding model-predicted effect.

Upon depicting the results of the analysis above, an unanticipated finding emerged: cTBS to SFS also had a negative effect on *Temporal Control*. A post-hoc 2 × 2 × 2 ANOVA examining effects of *Contextual Control*, *Temporal Control*, and *Target* comparing SFS to S1 revealed a similar pattern to that observed with cTBS to FPl: a significant *Temporal Control* × *Target* interaction in ER (F(1,21) = 8.99, p < 0.01), but not RT (F(1,21) < 1), and no interactions between *Contextual Control* × *Target* either alone or with *Temporal Control* (ER and RT, all p > 0.2). This result was not predicted by the model and no such effects were observed comparing cTBS to VLPFC with S1 (all interactions with *Target* p > 0.1). However, it is important to note that using DCM requires 2 steps: first is a model comparison step wherein plausible models of neural dynamics are compared; and second are inferences on the parameters of the best model of the data. Given that SFS did not exhibit a positive univariate effect of *Temporal Control* in our original study, we did not include SFS modulations as a function of *Temporal Control* in our original model. However, taking the previously estimated model as a starting point, the SFS could potentially influence *Temporal Control* through interactions with the mid-rostral LPFC (MFG) or FPl. Thus, a behavioral impairment on *Temporal Control* following cTBS to SFS could be observed if SFS positively modulates MFG or FPl to facilitate *Temporal Control*.

### Network Dynamics Revisited

To examine whether the observed impact of SFS on *Temporal Control* could be due to a previously unappreciated modulation of SFS on rostral LPFC, we initially compared the model we described previously (Figure 1; Figure 2 – figure supplement 3), to models in which the SFS modulates processing in either the MFG or FPl during *Temporal Control*. This initial comparison revealed that models containing SFS-MFG connectivity and modulations were a substantially better match to the data (family model exceedance probability 0.9992). We then proceeded to thoroughly explore the model space assuming fixed connectivity between the SFS and MFG, and varying the pathways by which modulations of connectivity supported cognitive control.

Group-level Bayesian model comparison adjudicated between models of effective connectivity within the LPFC (Stephan et al., 2009) (see Methods for full details of the procedure). Significant parameter estimates resulting from the revised best model are depicted in Figure 4 for both the previously described data (Figure 4 – figure supplement 1A), the present data (Figure 4 – figure supplement 1B), and the data considered jointly (Figure 4). The major difference from the original model is that the feedback modulation from FPl to MFG as a function of *Temporal Control* has been replaced by a feedforward modulation from SFS to MFG. There is also evidence of a correspondingly negative feedback modulation from MFG to SFS. In this model, disruption of SFS by cTBS would be predicted to impair *Temporal Control* by disrupting feedforward influences from SFS to MFG. This modulation was not necessitated by the model comparison or estimation procedures. Models containing no feedforward modulation from SFS to MFG were directly compared to models containing this influence in several steps of the model space exploration. Furthermore, the model comparison procedure only posits what modulations might exist, not the sign or strength of the modulation. Hence, even within models containing the feedforward influence from SFS to MFG, the modulation could have been negative rather than positive (or even non-significant). That the best model of the data contained this positive modulation from SFS to MFG is therefore in striking agreement with the behavioral effects of cTBS, demonstrating the utility of combining models of regional dynamics with neural perturbations (Jazayeri and Afraz, 2017).

**Figure 4:**
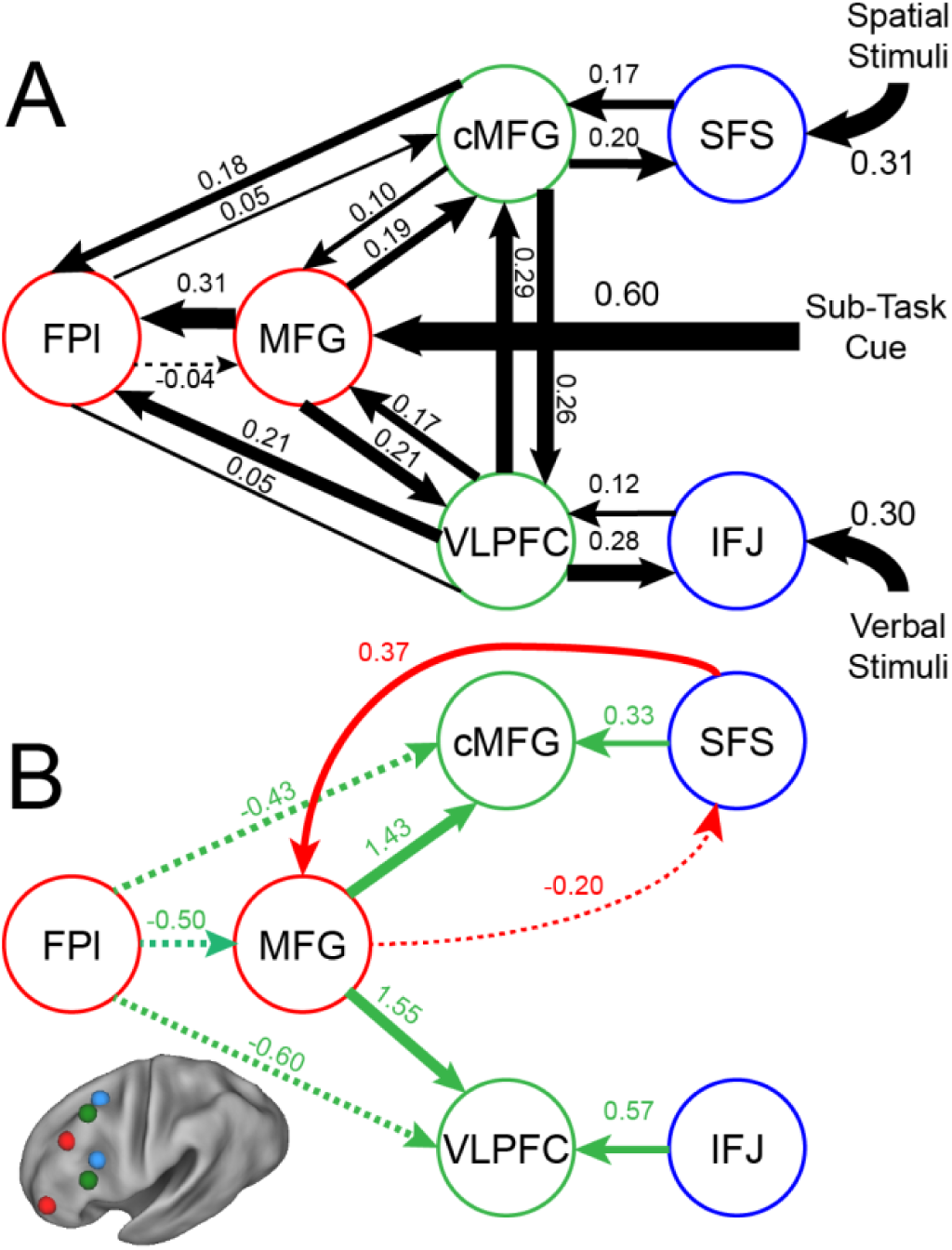
Revised LPFC dynamic causal model. Bayesian model selection indicated that the depicted model was the best model of the dynamics among the models tested. Arrows indicate direction of influence, numbers and line widths indicate the strength of influence, and dashed arrows indicated inhibitory influences. Parameter estimates have been averaged across the previous and present samples. A) Fixed connectivity and inputs depicted in black. B) Modulations of connectivity by *Contextual Control* (green), and *Temporal Control* (red) demands depicted in colors. All depicted parameters are significant after correction using false-discovery rate.

Finally, as in our original study, we examined measures of hierarchical strength and the relationship between model parameters and trait-measured higher-level cognitive ability. In the new model, effects of hierarchical strength remained highest in mid LPFC (both samples combined, positive peak of fitted vertex t(47) = 11.63, p < 10^-14^; mean position of vertex y = 21.7 in MNI space; Figure 4 – figure supplement 2). However, no relationship was observed between model parameters and trait-measured higher-level cognitive ability (both samples combined, hierarchical strength: r = -0.12, p > 0.4; top-down strength: r = 0.18, p > 0.2; bottom-up strength: r = 0.10, p > 0.4). While these data cast doubts on whether the model parameters can predict trait-measured cognitive ability, the good agreement between the effects of cTBS on behavior and the model predictions add validity to the model’s relationship to behavioral manifestations of cognitive control.

## Discussion

We used cTBS to test the causal relationship between a neural model of LPFC dynamics supporting cognitive control and behavioral manifestations of these control processes. In line with the model’s predictions, cTBS to caudal LPFC disrupted stimulus feature processing, cTBS to mid LPFC disrupted responding to a stimulus feature based on prevailing contextual contingencies (*Contextual Control*), and cTBS to rostral LPFC disrupted the ability to balance ongoing and future demands (*Temporal Control*). Unexpectedly, cTBS to caudal LPFC also disrupted *Temporal Control*. This result could not be accommodated by our previously modeled connectivity dynamics (Nee and D’Esposito, 2016). However, a revised model that included a modulation from caudal LPFC to mid-rostral LPFC provided a better fit to the neural data and a ready explanation for the observed impairment. Collectively, these results demonstrate the mutually informative nature of models of neural dynamics and causal brain-behavior tests, and reveal how the LPFC supports cognitive control.

cTBS is presumed to reduce cortical excitability (Huang et al., 2005) providing the ability to modulate behaviors for which neural activity of a stimulated region is critical. It is commonly assumed that applying cTBS to regions that show increased activity during a particular behavior will disrupt that behavior insofar as that area is necessary. With this logic in mind, it may be surprising to find that cTBS to caudal LPFC impacted *Temporal Control* since this area show *decreased* rather than increased activity during *Temporal Control* (previous/present sample: F(1,23) = 14.62/6.77, p’s < 0.05). cTBS presumably decreased activity in this region further, which is inconsistent with a simple mapping between univariate activity and behavior (i.e. further decreasing activity may have predicted a behavioral improvement). Moreover, the VLPFC showed increased activation during *Contextual Control* (previous/present sample: F(1,23) = 43.65/126.82, p’s < 10^-5^), and verbal processing (previous/present sample: F(1,23) = 25.8/16.05, p’s < 0.001), but not their interaction (previous/present sample: F(1,23) = 3.29/3.32, p’s > 0.05). Yet, cTBS to this region specifically impaired the interaction among these processes. That behavioral impairments following cTBS do not necessarily track simple regional increases/decreases in activity underscores the need to inform predictions of neural perturbation by models of neural dynamics.

Causal methods provide strong tests of hierarchical relationships. The logic of this approach can be understood by visualizing hierarchy as a pyramid – perturbation of the lowest levels of the hierarchy affect all levels above, while perturbation of the top level of the hierarchy does not disrupt the lower levels. The underlying neural mechanism of this hierarchical organization can be understood by the flow of information within hierarchical networks. Both transcranial magnetic stimulation (TMS) and lesion data have been used to examine the organization of the LPFC and hierarchical dependencies with varying results. Badre et al (2009) found that damage to caudal LPFC impaired the ability to base responses on color-feature associations (concrete, lower level deficit), while damage to mid LPFC impaired the ability to base responses on color-feature dimension associations (abstract, higher level deficit). These deficits were alternately described as hierarchical (higher level deficits were more likely to occur when lower level deficits were present), and independent (cross-over interaction between feature-caudal LPFC and dimension-mid LPFC deficits). It is unclear how to reconcile this dependence in behavioral levels, but independence among brain regions. One possible resolution is that a third region is necessary for the functions of both caudal and mid-LPFC that, when damaged, produced both lower and higher level deficits. Such a region could either provide feature inputs to caudal and mid-LPFC (e.g. SFS), or produce outputs that are informed by the processing/representations in caudal and mid-LPFC (e.g. premotor cortex). In the latter case, the predicted dynamics would be top-down modulations from caudal and mid-LPFC to premotor cortex during their respective forms of control, but not between one another. Such matters may have been difficult to resolve in the examined patient sample (11 patients). Resolving functional (in)dependence can also be difficult in brain-damaged patients whose lesions do not respect functional boundaries. Nevertheless, a model of neural dynamics on the studied task (Badre and D’Esposito, 2007) may help to inform these issues.

Rahnev et al (2016) and Bahlmann et al (2015) applied TMS to different frontal sites along the rostralcaudal axis and reported dissociable effects of stimulation, but no hierarchical dependencies. In both of these cases, lesions/stimulation were guided by examination of regional neural activity (i.e. univariate effects) rather than by models of neural dynamics among regions. These data suggest that under certain circumstances, frontal areas independently contribute to behavior. By contrast in a study of brain-damaged patients, Azuar et al (2014) reported hierarchical dependencies along the rostral-caudal axis in a task containing three levels of control: color-response mapping (low), color-task-response mapping (mid), episode-color-task-response mapping (high). In particular, patients with lesions to caudal LPFC were impaired at all forms of cognitive control, patients with lesions to mid LPFC were impaired at mid and high levels of control, while patients with lesions to rostral LPFC were impaired only at high levels of control. These data are consistent with a model of neural dynamics indicating a rostral-to-caudal cascade of activity whereby progressively rostral areas influence caudal areas as the level of cognitive control demands increases (Koechlin et al., 2003; Kouneiher et al., 2009). Collectively, these results indicate that whether or not hierarchical dependencies exist depends critically upon task-elicited regional interdependencies.

Our original report challenged prevailing accounts positing that the rostral LPFC (i.e. FPl) is the apex of the LPFC hierarchy. By definition, a higher place in a hierarchy entails greater influence over that which is lower than vice versa (Badre and D’Esposito, 2009). Previously, we found that mid areas of the LPFC showed the presumed signature of a hierarchical apex – greater outward than inward influence in fixed connectivity. This connectivity pattern was also observed in the present dataset, and in the revised model in both datasets. Furthermore, we previously observed that modulations of connectivity dynamics indicated that during cognitive control, activity converges towards mid LPFC. This result was also observed in the revised model, although some of the pathways by which information flows through the network have been altered. Nevertheless, it is clear that activity diverges (fixed connectivity) and converges (connectivity modulations) to/from the mid LPFC in the modeled neural dynamics. Collectively, these data corroborate the hypothesis we posited previously that the mid LPFC is a critical nexus for cognitive control. However, as indicated above, the criticality of a given region for cognitive control is task/demand dependent. Another test for the idea that the mid LPFC is a nexus would be to demonstrate the diversity of demands under which it is critical (Bertolero et al., 2015), indicating particular behaviors for which the mid LPFC is a dynamic hub (Osada et al., 2015). Examining the range of tasks affected by perturbation of different LPFC areas is a promising avenue to further examine the importance of different frontal regions for cognitive control more broadly.

While the significant effects observed in the present study were by-and-large predicted by a model of LPFC dynamics, the effects were generally limited by task accuracies that were near ceiling (average performance across the task ∼95%). Since error trials are typically considered separately in fMRI data, we included extensive practice to ensure that participants could perform the task with maximizing power for the fMRI data in mind. However, that many of the behavior modulations resulting from cTBS would be borne out in errors was not anticipated. A more suitable procedure may have been to titrate the task to individual performance levels, potentially by requiring adaptively speeded responses to keep performance from ceiling. Given these ceiling effects, it is difficult to interpret null effects in the cTBS data even though null effects are also important for validating the model. Future explorations using these methods would do well to keep these considerations in mind.

## Materials and Methods

Many of the materials and methods are identical to our previous report (Nee and D’Esposito, 2016). We present abbreviated details here, highlighting differences, as well as the cTBS procedures.

### Participants

We report results from 24 (15 female) right-handed native English speakers (mean age 20.5 years, range 18-27). Informed consent was obtained in accordance with the Committee for Protection of Human Subjects at the University of California, Berkeley.

The targeted number of participants was based upon previous work. A sample size of 24 participants was acquired to match to our previous study, which was well-powered to examine fMRI effects. We also considered the efficacy of cTBS. A previous study including a subset of the authors used a similar cTBS design with a smaller sample size of 17 (Rahnev et al., 2016) suggesting that our target sample would likely be adequate. Also, a sample size of 24 offered the ability to perfectly counter-balance the order of cTBS stimulation targets. All 24 participants are included for the fMRI analyses. In one participant, the control site (S1) was mistargeted, resulting in stimulation of an area that was active for the task (superior parietal lobule). The cTBS data from this participant is excluded. Another participant appeared to misunderstand the instructions for the *Restart* condition, demonstrating below chance accuracy on return trials. This participant was excluded for all cTBS analyses that included cognitive control demands, but was included in the analysis of *Stimulus Domain* with trials from the *Restart* condition excluded. Inclusion of these data was deemed appropriate over outright exclusion given that the analysis of *Stimulus Domain* was orthogonal to the cognitive control demands, and given the costs in data acquisition. These exclusions resulted in 23 participants for cTBS analyses focused solely on *Stimulus Domain*, and 22 participants for the remainder of the cTBS analyses.

Three participants performed the fMRI session but were excluded from cTBS sessions. For two participants, functional mapping revealed right-lateralized language processing despite self-reported right-handedness. Another participant was excluded due to an incidental finding. Three participants did not complete all of the cTBS sessions and were therefore excluded from analyses. Two of these participants experienced transient discomfort during stimulation of the rostral LPFC target, which mandated aborting that session and subsequent sessions. One participant withdrew from the study after the first cTBS session. These data were excluded from all analyses. Additionally, due to a technical error, the final 2 runs of data were not collected for one session for one participant.

### Procedure

The task design was a factorial 2 × 2 × 2, with factors of *Stimulus Domain* (verbal, spatial), *Contextual Control* (high, low), and *Temporal Control* (high, low). The procedure was nearly identical to our previous report and details can be found there (Nee and D’Esposito, 2016). Here we report differences in the procedure.

Whereas previously, participants performed 2 fMRI sessions, only a single fMRI session was performed here. The number of trials of the basic task at the beginning of each block was reduced from 2-5 trials to 2-3 trials, which helped to improve trial yield for trials-of-interest in the cTBS sessions. Otherwise, the fMRI session was identical to the previously described procedure.

Timing details were mistakenly omitted in our previous report. For fMRI sessions, stimuli were presented for 500 ms followed by a variable inter-trial interval (ITI) of 2600-3400 ms. Feedback indicating the number of correct trials in the block out of the total number of trials in the block was presented for 500 ms at the end of the block. Blocks were spaced by a variable 2600-3400 ms interval. Self-paced breaks were administered in-between runs. For cTBS sessions, the ITI was reduced to 1500-1900 ms, and the interval between blocks was reduced to 1100-1900 ms in order to improve trial yield. Breaks between runs were fixed at 30 s. With these timing reductions, 8 runs were administered resulting in 16 blocks for each cell of the design. It has been reported that reduced cortical excitability following cTBS begins approximately 5 minutes post-stimulation and lasts until approximately 60 minutes post-stimulation (Huang et al., 2005). Our timing procedures closely matched this window. A practice run of the task that was half the duration of the experimental runs immediately followed cTBS. As a result, experimental runs began approximately 5 minutes post-stimulation. A preliminary cursory inspection of time effects (indexed by run within session) revealed no appreciable patterns, so effects of time were not further explored.

Participants performed six sessions scheduled as follows: participants first completed a behavioral session wherein they were introduced to the task and given extensive practice as previously described. Assays of higher-level cognitive ability were also collected as previously described. The fMRI session followed on a separate day, within 10 days of the behavioral session (1 participant was scanned 17 days after the behavioral session due to a scheduling issue). The first cTBS session followed the fMRI session within 10 days (12 days for one participant with a scheduling issue). All cTBS sessions were spaced by one week with each session beginning at the same time of day for a given individual. The order of stimulation targets was fully counterbalanced and randomized across participants.

### fMRI Analysis

Analysis of the fMRI data was virtually identical to the methods previously described (Nee and D’Esposito, 2016). Regions-of-interest (ROI) analyses were centered on peak coordinates of our previous report. The one exception to this was for the caudal middle frontal gyrus (cMFG) which was centered on peak activation for *Contextual Control* in the present sample (-26 14 52) to accommodate individual variability in slice prescription that left out the previous peak (-34 10 60) in some participants. ROIs were used to explore abstraction effects both by analysis of *Stimulus Domain* and by correlations with behavior, as previously described.

### Transcranial Magnetic Stimulation Procedures

Transcranial magnetic stimulation (TMS) was delivered using a MagStim Super Rapid2 stimulator equipped with a figure-eight double air film coil with a 70 mm diameter. Electromyography (EMG) was recorded using electrodes place on the first dorsal interosseus (FDI) muscle on the right hand using a Delsys Bagnoli system (Delsys Inc.). Individual active motor threshold (AMT) was determined immediately before each session of cTBS. First, a “hunting procedure” was used to determine the scalp location in the left hemisphere producing the maximal contralateral hand twitch at the minimal stimulation intensity. Next, the participant maintained voluntary contraction of the FDI muscle at ∼20% of maximum contraction. AMT was determined as the minimal stimulation intensity needed to produce a transient cessation of EMG activity in 5 out of 10 pulses. Across sessions and participants, the average AMT was 48% of the stimulator output.

cTBS was delivered in a standard manner as described by (Huang et al., 2005). Bursts of three pulses at 50 Hz were delivered every 5 Hz for a total of 600 pulses over 40 s. Stimulation intensity was delivered at 80% of the individual’s AMT. During all TMS procedures, participants were continually monitored and verbally queried for signs of pain, dizziness, or other adverse effects. Just prior to the start of cTBS, a single test pulse was delivered to the target site at the same intensity of cTBS. The test pulse was used to help the participant determine whether stimulation of the target would be painful. As reported above, two participants found stimulation of rostral LPFC painful during this test and were excluded from further participation. No adverse effects were reported in the sample used for analysis.

### cTBS Targets

cTBS targets were defined based upon individual activation maxima (LPFC) or anatomy (S1). We restricted targets to the left hemisphere. LPFC targets were chosen to dissociably stimulate caudal, mid, and rostral LPFC. As reported previously, both dorsal and ventral peaks exist along the rostral-caudal axis of the LPFC. The choice of which target to stimulate for each rostral/caudal sector was motivated by the desire to maximize the distance between targets. Furthermore, given that the ventral caudal LPFC area (i.e inferior frontal junction; IFJ) was distanced from the scalp surface (mean MNI peak -38 6 26 previously), the dorsal caudal LPFC was selected as a more suitable target. This lead to targeting the ventral mid LPFC and FPl rostrally.

The caudal LPFC target was chosen as the maximal activation for the *Stimulus Domain* contrast whose sign was dictated by the contrast of *Spatial Stimulus Domain* > *Verbal Stimulus Domain* (mean MNI location: -23.9 -0.6 55.6). Previously, it was found that the mid LPFC shows univariate sensitivity to both *Stimulus Domain* and *Contextual Control*, but not their interaction. As can be observed in the whole-brain univariate maps (e.g. Figure 2), the *Contextual Control* contrast elicits activation along much of the rostral-caudal axis of the LPFC potentially rendering identification of the appropriate stimulation target difficult on an individual basis. We thus chose to use the *Stimulus Domain* contrast to define the mid LPFC target since this contrast produced more confined activations that were anticipated to reduce variability in target location across participants. In this case, areas were defined as those showing greater activity for the *Verbal Stimulus Domain* > *Spatial Stimulus Domain*. The ventral mid LPFC area previously described extended from the inferior frontal sulcus (IFS) into the inferior frontal gyrus ventrally, and middle frontal gyrus dorsally. To maintain consistency, we chose targets restricted to the inferior frontal gyrus. As a result, we now describe the ventral mid LPFC area as the ventrolateral prefrontal cortex (VLPFC) since this term more accurately describes the targeted site. For the first 6 participants, the maximal activation on the lateral surface of the inferior frontal gyrus was chosen as the stimulation target. At this point, it was determined that this choice of target definition was producing targets that were consistently caudal to the ventral mid LPFC area previously described (mean MNI y-coordinate of the first 6 participants = 14.0 compared to 20 previously). The procedure was therefore adjusted to target the rostral-most local maxima within pars triangularis and pars opercularis. This resulted in targets that were more clearly situated in mid LPFC (mean MNI location: -51.3 17.5 17.5).

The rostral LPFC target was chosen to be in the vicinity of the FPl previously described (mean MNI location: -34.1 56.9 0.7). In most cases, the rostral LPFC target was identified as the maximal activation for the conjunction of *Temporal Control* and *Contextual Control*. However, if the activation peak was deemed too close to the orbits to stimulate without pain, a more dorsal local maxima was used. In 5 cases, a target could not be identified through the conjunction contrast. In these cases, either the main effect of *Temporal Control*, main effect of *Contextual Control*, or the simple contrast of Branch > Control was used with the main effect contrasts taking precedence. This procedure resulted in a target that, on average, was equivalently engaged during *Temporal Control* and *Contextual Control* (average t-statistic for *Temporal Control*: 4.15, average t-statistic for *Contextual Control*: 4.38). Finally, the control site (S1) was chosen as the dorsal-most portion of the post-central gyrus anatomically defined. cTBS targets were guided by a frameless stereotactic localization system (Brainsight; Rogue Research, Inc.).

### cTBS Analysis

Behavioral modulations following cTBS were analyzed using repeated measures ANOVAs. For the analysis of *Stimulus Domain*, all trials were considered in order to maximize power. This was because the demand to select a stimulus feature (either verbal or spatial) was present in all trials of the task with the potential exception of the sub-task phase of *Delay* trials (notably, removing Delay trials from the analyses does not change the results). Other analyses were restricted to trials from the sub-task phase wherein the other cognitive control demands were manipulated. Reaction times (RTs) were computed from correct trials only. RTs greater than 2.5 standard deviations of the condition mean were discarded as outliers, as were trials with RTs less than 200 ms or greater than 2000 ms. ∼3.5% of the trials were removed as outliers. For analyses on accuracy/error-rate, condition-wise measurements of % correct were arc-sine transformed to normalize the distributions. Accuracies at ceiling (i.e. 100% correct) were corrected prior to transformation with the formula (*n*-0.5)/*n* where *n* is the number of trials in the condition. Statistics were performed on these transformed data. To present error-rates, transformed accuracies were subtracted from the corrected transformed ceiling. Error-rates are presented rather than accuracies to facilitate interpretation alongside RTs such that greater numbers consistently reflect worse performance.

To test for contrast × Target interactions, the contrasts of the main effect of *Stimulus Domain* in ER, *Stimulus Domain* × *Contextual Control* in RT, and *Temporal Control* in ER were computed and z-scored across individuals. These were entered into a 3 × 4 repeated measures ANOVA. A follow-up 4 × 4 ANOVA added the contrast of *Contextual Control* in ER.

### Dynamic Causal Modeling

DCM was originally performed as previously described (Nee and D’Esposito, 2016). The presence/absence of connectivity pathways and modulations was matched to our previous report and the model was estimated in the present sample. Inferences on the parameter estimates were performed in an identical manner as described previously.

Given the unanticipated effect of cTBS to caudal LPFC on *Temporal Control*, additional DCM analyses were performed on the present sample. First, the original model was compared to two other families of models: 1) Fixed connectivity between the SFS and FPl along with modulations (either feedforward, feedback, or bi-directional) as a function of *Temporal Control*, and 2) Fixed connectivity between the SFS and MFG along with modulations (either feedforward, feedback, or bi-directional) as a function of *Temporal Control*. Bayesian model selection was first performed family-wise (Penny et al., 2010). In this comparison, families including connectivity between SFS and MFG were overwhelmingly favored (model exceedance probability = 0.9992). Given this result, all subsequent model comparisons included fixed connectivity between the SFS and MFG, with modulations as a function of *Temporal Control* (feedforward, feedback, bi-directional, or absent) varying across models. We used an iterative procedure as previously described to explore the model space while maintaining computational feasibility (Nee and D’Esposito, 2016). Here, the model space was identical to before with the addition of the possibility of modulations between the SFS and MFG as a function of *Temporal Control*. In this case, the iterative procedure did not converge, instead hitting an infinite loop. In total, 913 models were explored. Simultaneous comparison of these models using Bayesian model selection (Stephan et al., 2009) did not identify a single best model in terms of an exceptional model exceedance probability. A scree plot of the models ordered by their exceedance probabilities identified potential step-wise differences for the top 18 and top 27 models. It is possible that the failure to identify a clearly superior model was due to the large number of models explored that overlapped in many features. Hence, Bayesian model selection was performed on a reduced model space taking the top 27 models. Once again, no single model was clearly favored, but the top 19 models (all model exceedance probability ∼0.04) were ordered similarly to before and were once again consistently favored over the other models (model exceedance probabilities ∼0.02). Nevertheless, these model exceedance probabilities were no different than uniform (1/27 = 0.037).

In order to better adjudicate between the models of LPFC dynamics, the best 19 models identified above were compared within the previous sample. Given that the previous sample contained twice the fMRI data per subject, this offered the ability to produce more stable model parameter estimates with better potential to delineate among the models. The original model was also included for comparison to verify that the addition of SFS to MFG connectivity was prudent making a total of 20 models for comparison. In this case, far more convincing evidence was found for the top model (model exceedance probability = 0.2526, compare to a uniform of 0.05). Other models with better than uniform model exceedance probabilities shared the main features found significant in the data reported in the main text (feedforward modulation from SFS to MFG for *Temporal Control*, feedback modulation from FPl to MFG and MFG to cMFG/VLPFC for *Contextual Control*) bolstering the credibility of these effects. Parameter estimates for the best model were examined for the previous dataset and the present dataset and compared for consistency. Noting good agreement among the parameter estimates (Figure 4 – figure supplement 1), the two datasets were combined for subsequent inference. Random effects inference on the parameters of the winning model then proceeded in a manner identical to our previous report.

## Acknowledgements

This research was supported by National Institute of Neurological Disorders and Stroke Grant F32 NS0802069 (DN) and P01 NS040813 (MD), and National Institute of Mental Health Grant R01 MH063901 (MD). The authors thank Dobromir Rahnev for helpful advice with the TMS procedure, and Max Wang, Lara Krisst, Grace Lee, Joanne Lin, and Chelsea Harmon for help with data collection.

---

**Figure 2 – figure supplement 1:**
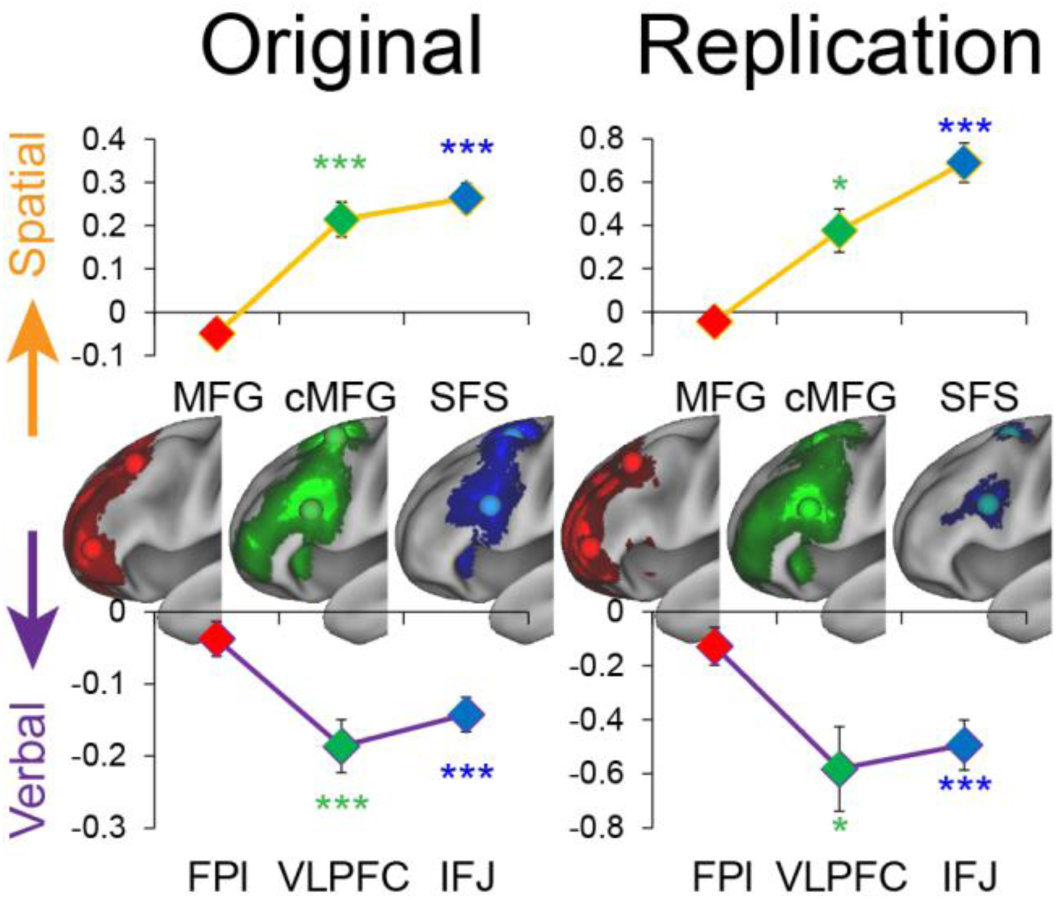
Stimulus domain abstraction replication. Left: previously reported univariate effects of *Stimulus Domain*. Right: Univariate effects of *Stimulus Domain* in the present study. The y-axis depicts the contrast of *Stimulus Domain* with positive effects indicating increasing spatial sensitivity and negative effects indicating increasing verbal sensitivity. Effects of *Stimulus Domain* were present in caudal (SFS: t(23) = 10.50, p_corrected_ < 10^-8^; IFJ: t(23) = -4.73, p_corrected_ < 0.001) and mid LPFC (cMFG: t(23) = 3.35, p_corrected_ < 0.05; VLPFC: t(23) = -3.70; p_corrected_ < 0.01), but not rostral LPFC (MFG: t(23) = -0.70, p > 0.4; FPl: t(23) = -0.87, p > 0.3) consistent with rostral areas performing abstract cognitive control that does not depend on stimulus features. Statistics reflect Bonferroni-corrected tests in the present study. * - p_corrected_ < 0.05; ** - p_corrected_ < 0.005; *** - p_corrected_ < 0.0005.

**Figure 2 – figure supplement 2:**
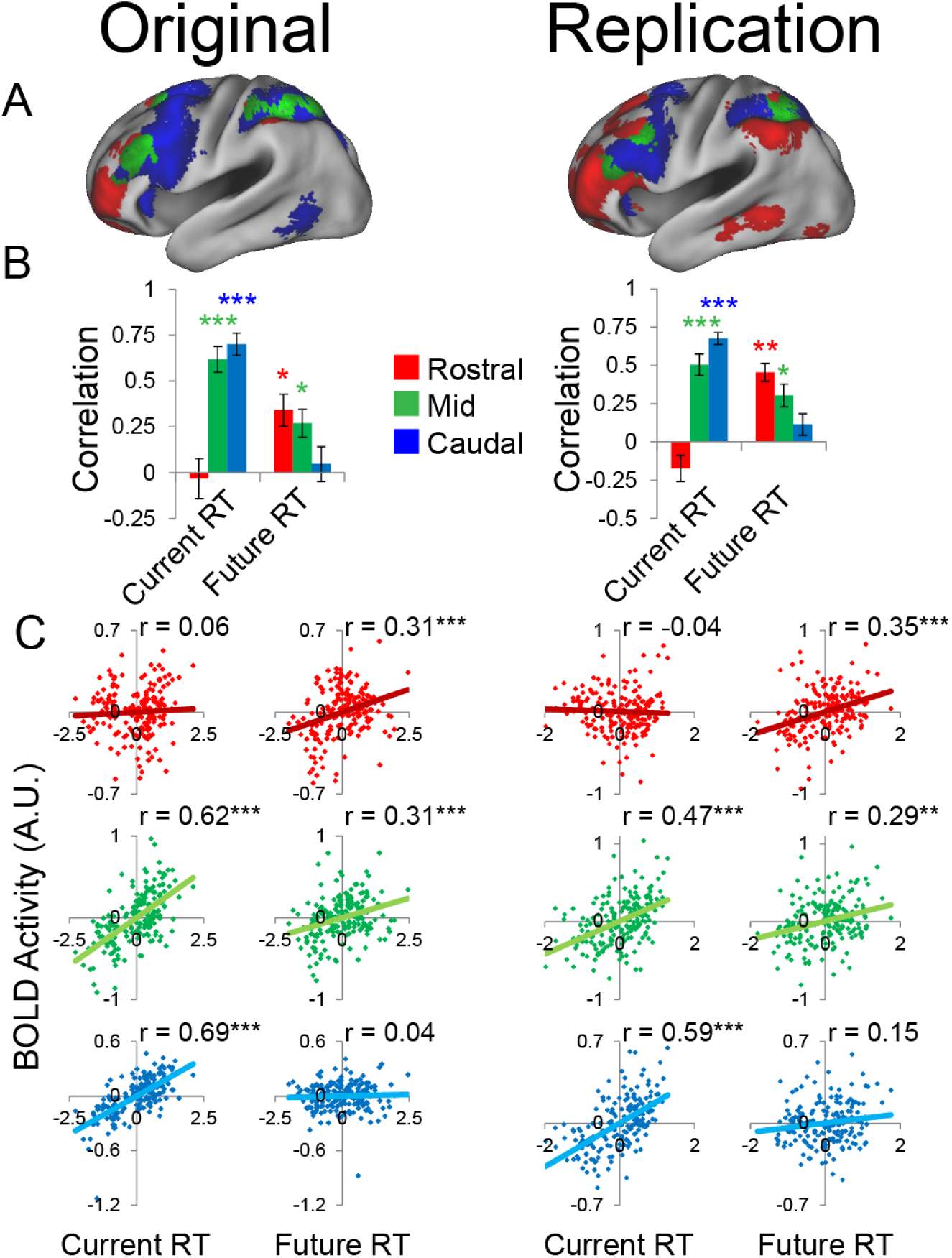
Temporal activation-behavior relationship replication. Left: previously reported partial correlations between activation and behavior. Right: Partial correlations between activation and behavior in the present study. Current RT corresponds to sub-task trials while future RT corresponds to return trials. RT measures have been normalized within-subject across the 8 conditions of interest. A) Voxel-wise regression of Current and Future RT onto activations for the 8 conditions of interest across subjects. Individual subject terms have been regressed out. Red: significant correlations with Future RT only; Blue: significant correlations with Current RT only; Green: both. B) Average partial correlation between activation and RT for the 8 conditions of interest computed separately for each subject (summary-statistic approach). C) Partial correlations between activation and RT for the 8 conditions of interest for all subjects. Individual subject terms have been regressed out. Red: rostral LPFC; Green: mid LPFC; Blue: caudal LPFC. In all cases, caudal LPFC was sensitive to Current (summary statistic approach: t(23) = 8.70, p_corrected_ < 10^-7^; full partial correlation: r = 0.59, p_corrected_ < 10^-15^), but not Future RT (summary statistic approach: t(23) = 0.74, p > 0.4; full partial correlation: r = 0.15, p_corrected_ > 0.3), rostral LPFC was sensitive to Future (summary statistic approach: t(23) = 4.29, p_corrected_ < 0.005; full partial correlation: r = 0.35, p_corrected_ < 0.0001), but not Current RT (summary statistic approach: t(23) = t(23) = -1.29, p > 0.2; full partial correlation: r = -0.04, p > 0.6), and mid LPFC was sensitive to both Current (summary statistic approach: t(23) = 5.36, p_corrected_ < 0.0005; full partial correlation: 0.47, p_corrected_ < 10^-9^) and Future RT (summary statistic approach: t(23) = 3.48, p_corrected_ < 0.05; full partial correlation: r = 0.29, p < 0.001). Collectively, these results indicate a temporal abstraction gradient. Statistics reflect Bonferroni-corrected tests in the present study. * - p_corrected_ < 0.05; ** - p_corrected_ < 0.005; *** - p_corrected_ < 0.0005.

**Figure 2 – figure supplement 3:**
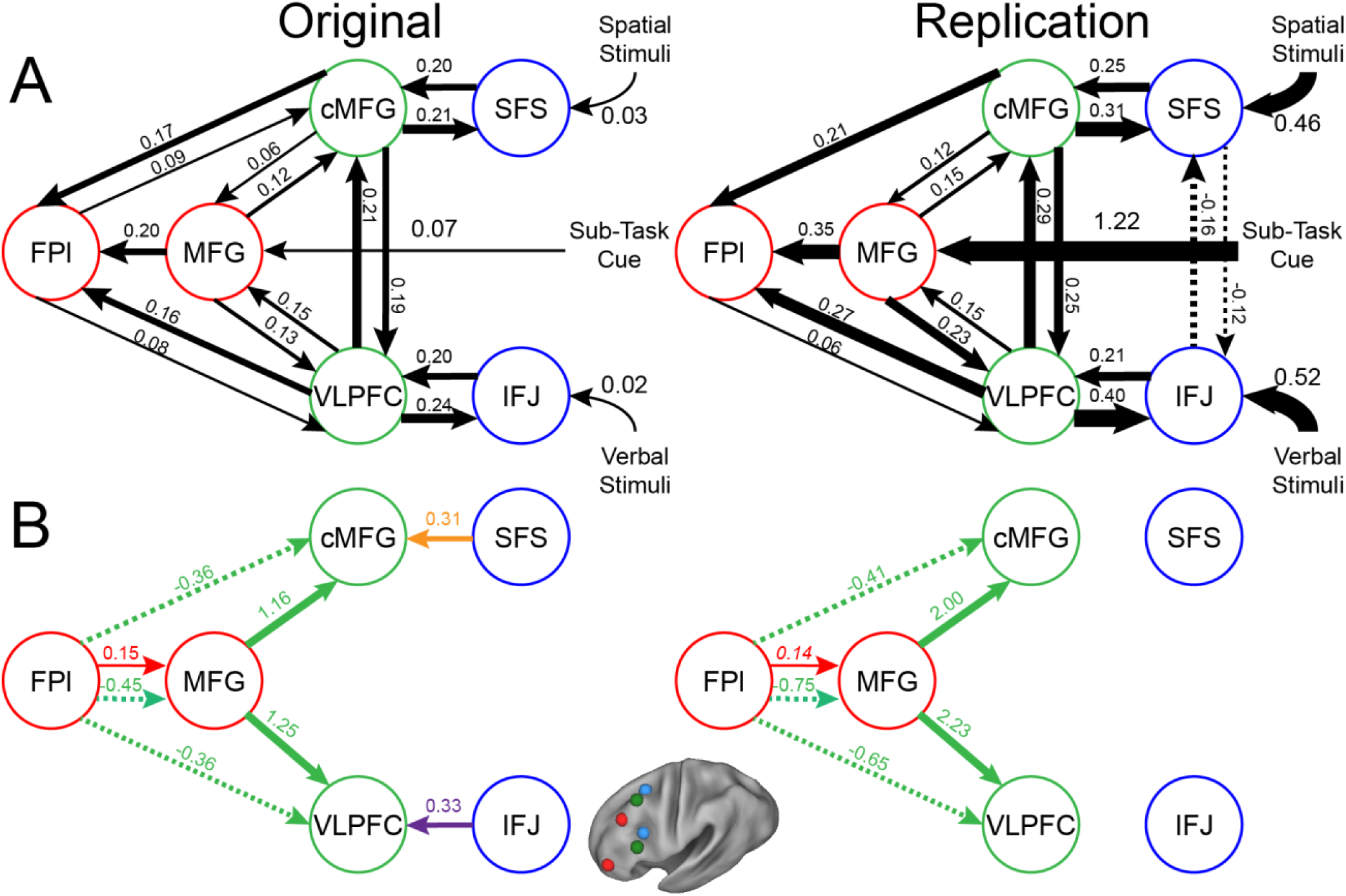
LPFC dynamic causal model replication. Left: parameter estimates for the best model indicated by random effects Bayesian model selection previously. Right: parameter estimates for the same model in the present data. Arrows indicate direction of influence, numbers and line widths indicate the strength of influence, and dashed arrows indicated inhibitory influences. A) Fixed connectivity and inputs depicted in black. B) Modulations of connectivity by *Spatial Stimulus Domain* (orange), *Verbal Stimulus Domain* (purple), *Contextual Control* (green), and *Temporal Control* (red) demands depicted in colors. All depicted parameters are significant after correction using false-discovery rate with the exception of the italicized modulation of connectivity between FPl and MFG in the replication sample which just missed significance (q = 0.057).

**Figure 2 – figure supplement 4:**
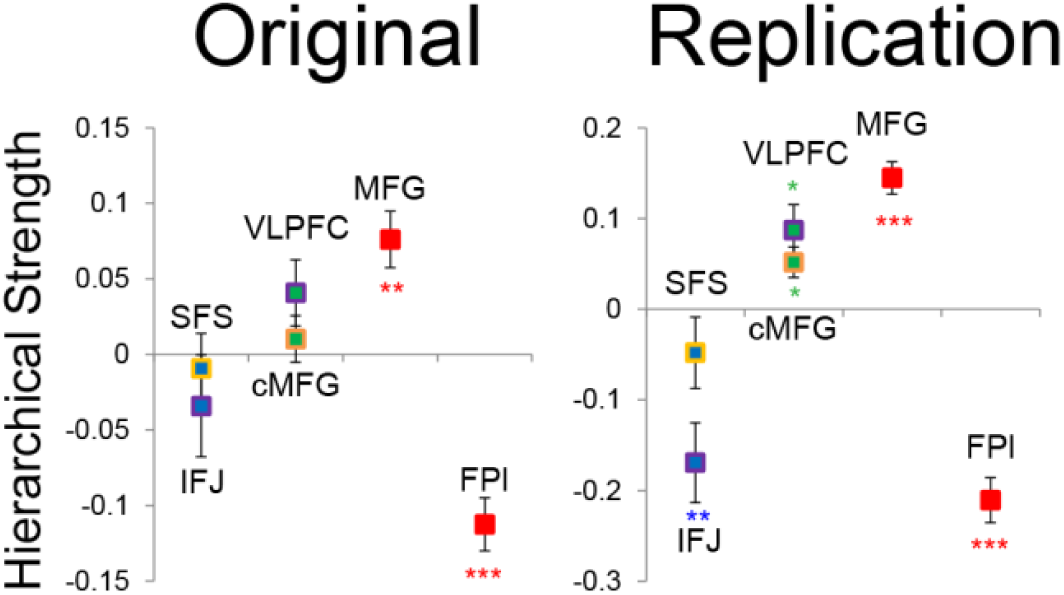
Hierarchical fixed dependencies replication. Based on fixed connectivity of the dynamic causal model, hierarchical strength was calculated as the difference between outward and inward projections along the rostral/caudal axis with relatively greater outward connectivity indicating more hierarchical strength. Left: previous data. Right: present data. A parabolic function fitting the relationship between rostral/caudal position (y-coordinate in MNI space) and hierarchical strength indicated a positive vertex across individuals (t(23) = 3.95, p < 0.001) positioned at mid LPFC (mean y-coordinate 27.5). ** - p_corrected_ < 0.005; *** - p_corrected_ < 0.0005.

**Figure 2 – figure supplement 5:**
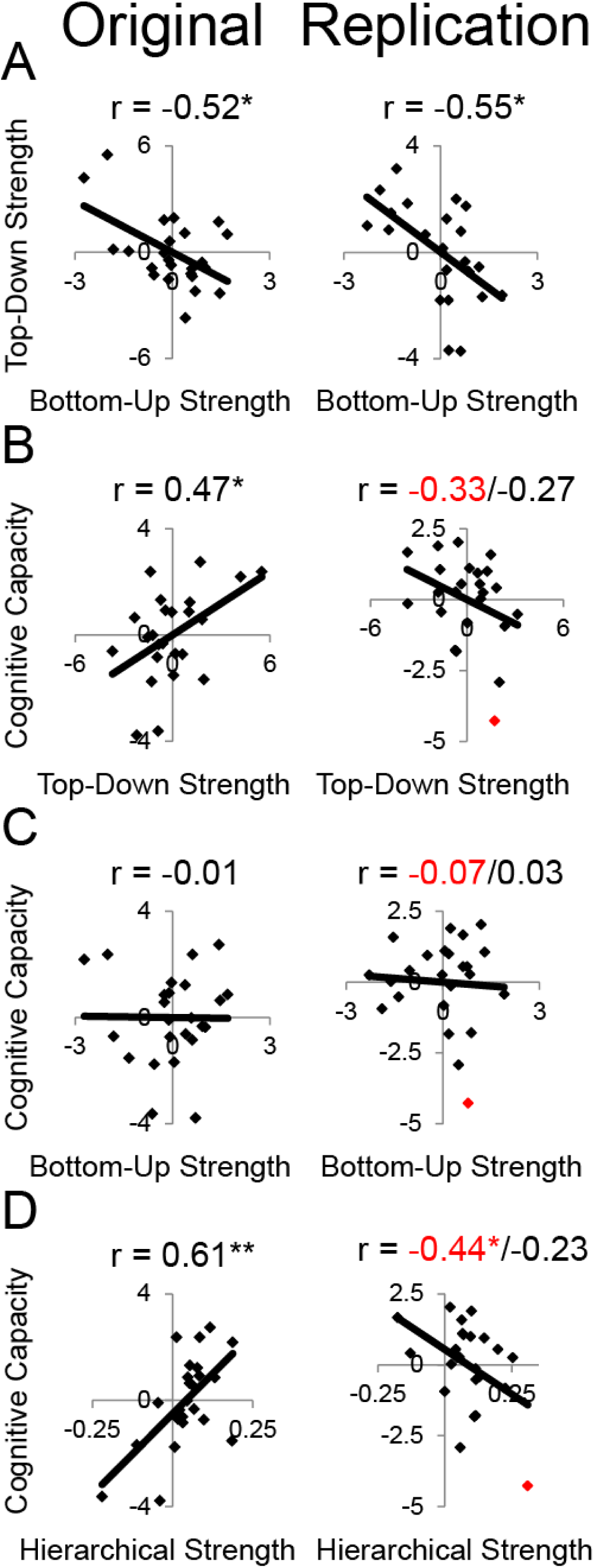
LPFC dynamics and higher-level cognitive ability replication failure. Neural metrics were based on modeled estimates of effective connectivity and their modulations. Metrics reflecting top-down LPFC modulations by cognitive control demands (top-down strength), and metrics reflecting bottom-up LPFC modulations by *Stimulus Domain* demands (bottom-up strength) were combined, respectively using principle components analysis. Left: previous data. Right: present data. A) Top-down and bottom-up strength were anticorrelated. B) Top-down strength predicted trait-measured higher-level cognitive capacity in the previous sample, but this pattern was non-significantly reversed in the present sample. Red indicates an outlier whose cognitive capacity was more than 2.5 standard deviations from the mean. With outlier included r = -0.33, p > 0.1; excluded r = -0.27, p > 0.2. C) Bottom-up strength did not correlate with higher-level cognitive capacity in either sample. D) Hierarchical strength reflected the degree to which mid LPFC showed greater outward than inward fixed connectivity. This metric was also positively related to higher-level cognitive capacity previously, but negatively in the present sample with an outlier included (r = -0.43, p < 0.05), but not with the outlier excluded (r = -0.23, p > 0.25). * - p < 0.05; ** - p < 0.005.

**Figure 3 – figure supplement 1:**
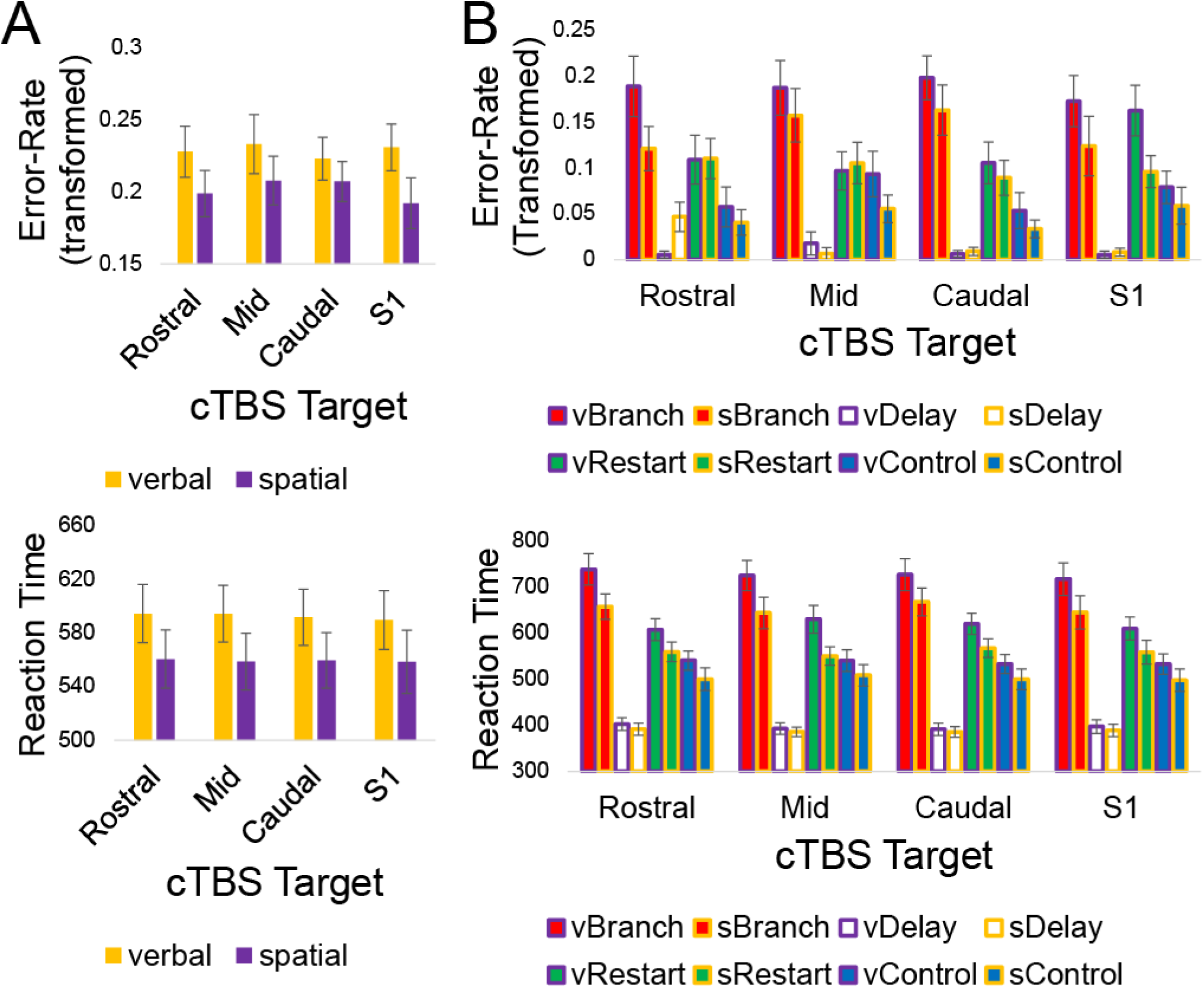
Full behavioral data. All of the behavioral data are depicted without contrasts.

**Figure 4 – figure supplement 1:**
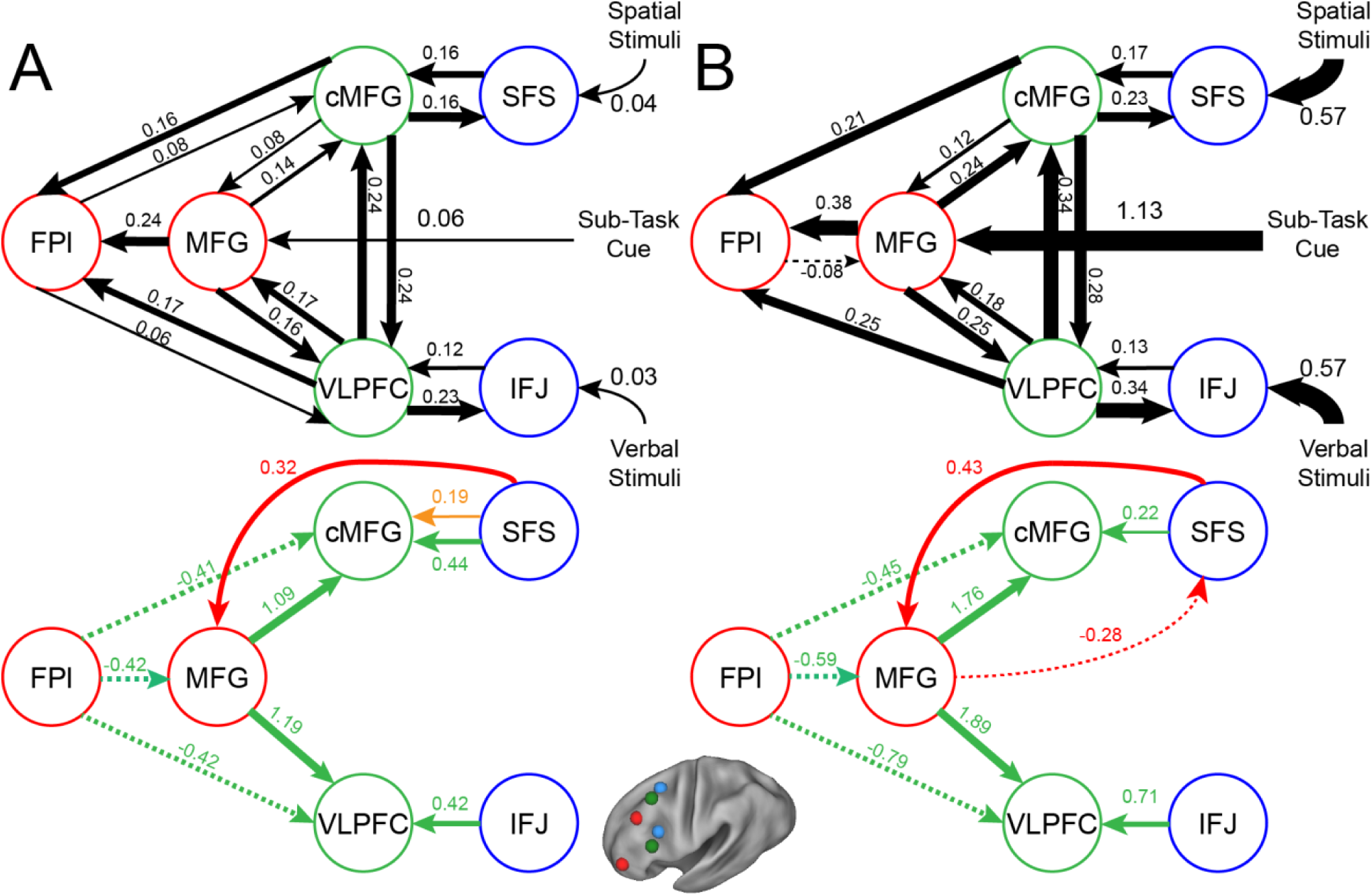
Revised LPFC dynamic causal model. Left: Original study data. Right: Current study data.

**Figure 4 – figure supplement 2:**
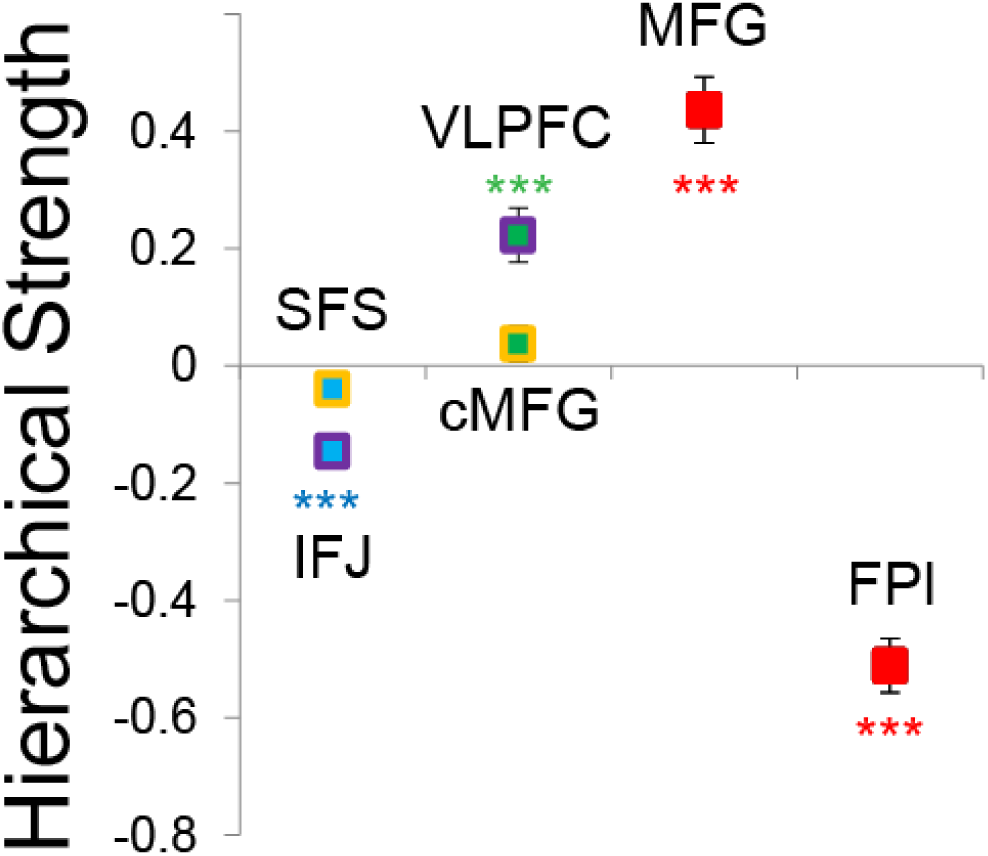
Revised hierarchical fixed dependencies. Hierarchical strength is detailed averaged across the previous and present samples. Details are otherwise identical to Figure 2 – figure supplement 4.

